# Fluorescence labeling strategies for the study of ion channel and receptor cell surface expression: A comprehensive toolkit for extracellular labeling of TRPV1

**DOI:** 10.1101/2024.05.09.593209

**Authors:** Taylor M. Mott, Grace C. Wulffraat, Alex J. Eddins, Ryan A. Mehl, Eric N. Senning

## Abstract

Regulation of ion channel expression on the plasma membrane is a major determinant of neuronal excitability, and identifying the underlying mechanisms of this expression is critical to our understanding of neurons. A critical aspect of measuring changes in ion channel expression is uniquely identifying ion channels located on the cell surface. To accomplish this goal we demonstrate two orthogonal strategies to label extracellular sites of the ion channel TRPV1 that minimally perturb the function of the channel: 1) We use the amber codon suppression technique to introduce a non-canonical amino acid (ncAA) with tetrazine click chemistry compatible with a trans-cyclooctene coupled fluorescent dye. 2) By inserting the circularly permutated HaloTag (cpHaloTag) in an extracellular loop of TRPV1, we incorporate a click-chemistry site for a chloroalkane-linked fluorescent dye of our choosing. Optimization of ncAA insertion sites was accomplished by screening residue positions between the S1 and S2 transmembrane domains with elevated missense variants in the human population, and we identified T468 as a rapid labeling site (∼5 minutes) based on functional as well as biochemical assays in HEK293T/17 cells. After several rounds of adapting the linker lengths and backbone placement of cpHaloTag on the extracellular side of TRPV1, our efforts led to a channel construct that robustly expressed as a fully functional TRPV1exCellHalo fusion with intact wild-type gating properties. The TRPV1exCellHalo construct was used in a single molecule experiment to track TRPV1 on the cell surface and validate studies that show decreased mobility of the channel upon activation. The success of these extracellular label TRPV1 (exCellTRPV1) constructs as tools to track surface expression of the channel will shed significant light on the mechanisms regulating expression and provide a general scheme to introduce similar modifications to other cell surface receptors.

## Introduction

The non-selective cation channel TRPV1 is expressed in the small-diameter class of dorsal root ganglia neurons known as nociceptive c-fibers (Bevan and Szolcsányi, 1990; Caterina et al., 1997). Activation of TRPV1 in these nociceptors by temperatures above 43 deg C is linked to physiological heat sensation, and both oxidation and acidity acting on TRPV1 lead to perception of noxious stimuli (Davis et al., 2000; Leffler et al., 2006; Chuang and Lin, 2009). Recent studies offer compelling evidence that TRPV1 function is central to inflammatory pain and, as such, has become a pharmacological target for pain intervention (Szolcsányi and Sándor, 2012).

As an important receptor for pain studies, TRPV1 channel research has made extensive progress along several aspects of its regulation, including desensitization by Ca^2+^, post-translational modifications, lipid regulation, and trafficking (Koplas et al., 1997; Olah et al., 2001; Morenilla-Palao et al., 2004; Zhang et al., 2005; Vetter et al., 2008). This is neither a complete list of TRPV1 mechanisms of regulation nor are the listed attributes necessarily mutually exclusive. In fact, the lipid PI(3,4,5)P3 is implicated as an essential signal for increased surface trafficking of TRPV1 (Stein et al., 2006; Stratiievska et al., 2018). Compelling evidence that surface expression of TRPV1 is an important aspect of the channel’s regulation has motivated studies that monitor trafficking of the channel (Morenilla-Palao et al., 2004; Zhang et al., 2005). Although voltage-clamp electrophysiology directly measures currents elicited from the total number of TRPV1 on the cell surface, there is an inherent ambiguity between current changes stemming from differences in single channel conductance versus differences in the total number of channels. As a complementary tool to electrophysiology, fluorescence imaging of cell surface channels and receptors fused with genetically encoded fluorescent proteins (FPs) is a bona fide approach to directly measure their changing numbers (Michaluk and Rusakov, 2022). By fusing a genetically encoded FP to the C-terminus of TRPV1, researchers can track the sub-cellular localization of the channel. Studies of TRPV1-FP surface expression in F11 DRG hybridoma cells and HEK293T/17 cells have informed our understanding about NGF dependent trafficking and Ca^2+^-dependent surface mobility of TRPV1(Stein et al., 2006; Senning and Gordon, 2015).

Two critical obstacles stand in the way of obtaining a more detailed mechanism behind TRPV1 trafficking and surface expression dynamics. The primary concern is that signal from fluorescently labeled channels on the surface must be discriminated from intracellular pools of the channel. If both populations of the channel are labeled with the same fluorescent protein, the signal to noise in the experiment is considerably less than if only the surface population were optically active. A secondary concern is that the fluorescent protein introduces a perturbation to the channel structure that may restrict aspects of its regulation(Wang et al., 2019). Moreover, channel function and interaction surfaces may be impacted by the inclusion of a large fluorescent protein domain.

Here we present a comprehensive set of fluorescence research tools that are intended to address the limitations of current TRPV1 labeling strategies, especially as concerns trafficking and sub-cellular localization. Our fluorescent labeling strategies rely on a post-expression cell surface click-chemistry approach with cell impermeable fluorescent chemical ligands (Figure 1A). We have optimized expression of a TRPV1-‘tag’ construct with an amber stop codon (‘tag’) to introduce an amino acid sized click-chemistry site with amber codon suppression techniques (Figure 1B,i)(Zagotta et al., 2016; Blizzard et al., 2018; Bessa-Neto et al., 2021). Optimal placement of our ‘tag’ site in the TRPV1 sequence was determined by genomic variant scanning, which builds on our previous work to relate variant positions identified in gnomAD with TRPV1 function (Karczewski et al., 2020; Mott et al., 2023). An alternative approach to extracellular labeling of TRPV1 was explored with the HaloTag click-chemistry system. We incorporated the circularly permuted HaloTag within an extracellular loop of TRPV1 (Figure 1B, ii)(Deo et al., 2021). Although the TRPV1exCellHalo construct sustains a large increase in molecular mass, we show that the channel expresses well, functions similarly to wild-type TRPV1, and assembles into heteromeric channels with C-terminally labeled TRPV1 subunits.

**Figure 1.**
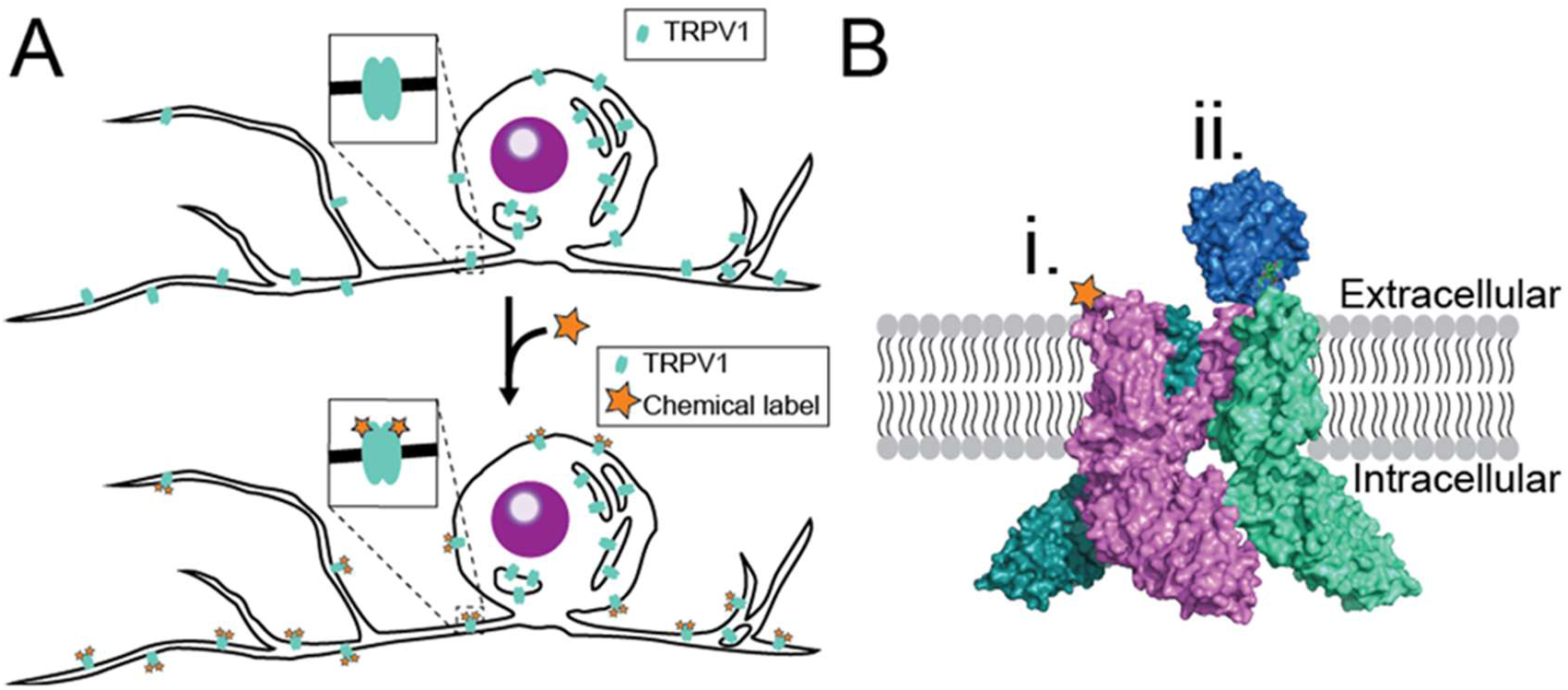
Two strategies for extracellular labeling of TRPV1. (A) TRPV1 expression in DRG neurons is distributed across the plasma membrane and intracellular compartments (top). If the extracellular side of TRPV1 can be labeled at a click-chemistry site, labeling the plasma membrane population of TRPV1 is possible (bottom). (B) Amber codon suppression techniques permit substitution of a non-canonical amino acid (ncAA) at a single residue in the TRPV1 sequence. Addition of an ncAA with click-chemistry properties in an extracellular loop of TRPV1 allows surface labeling of the ion channel (PDB 7lqy; B,i). By introducing the HaloTag domain (blue PDB 6u32) to an extracellular loop of TRPV1, cell impermeable HaloLigands can be specifically targeted to the population of channels that are expressed on the plasma membrane (B, ii).

## Methods

### Bioinformatic analysis of human genomic data

To examine the TRPV1 sequence for variant amino acid positions we accessed human genomic variant data from gnomAD 2.1.1. Human variant data for TRPV1 was downloaded directly from gnomAD servers (http://gnomad.broadinstitute.org; ‘‘export variants to CSV’’ option). Spreadsheet data was formatted in Excel (Microsoft) to read in the variant data to a Matlab (Mathworks, MA) script as done in Mott and coworkers that generated a frequency plot for the number of missense variants at each residue position between TRPV1 positions 427-680 (See Figure 2A) (Mott et al., 2023). Synonymous mutations are ignored, and each occurrence of a missense variant is recorded as a variant in our counts. Residues with >100 variants were truncated to 100 counts, and positional frequency was also smoothed with a boxcar average of three residues. Alignment with secondary structure was done with respect to rat TRPV1 residue positions, and alpha helical structure was based on the TRPV1 secondary structure assignment in Nadezhdin and coworkers (Nadezhdin et al., 2021). Sequence alignment data for conservation analysis was obtained from Uniprot.org (www.uniprot.org) and alignments are displayed in Figure 2D with alignment annotator (http://bioinformatics.org/strap/aa/)

**Figure 2.**
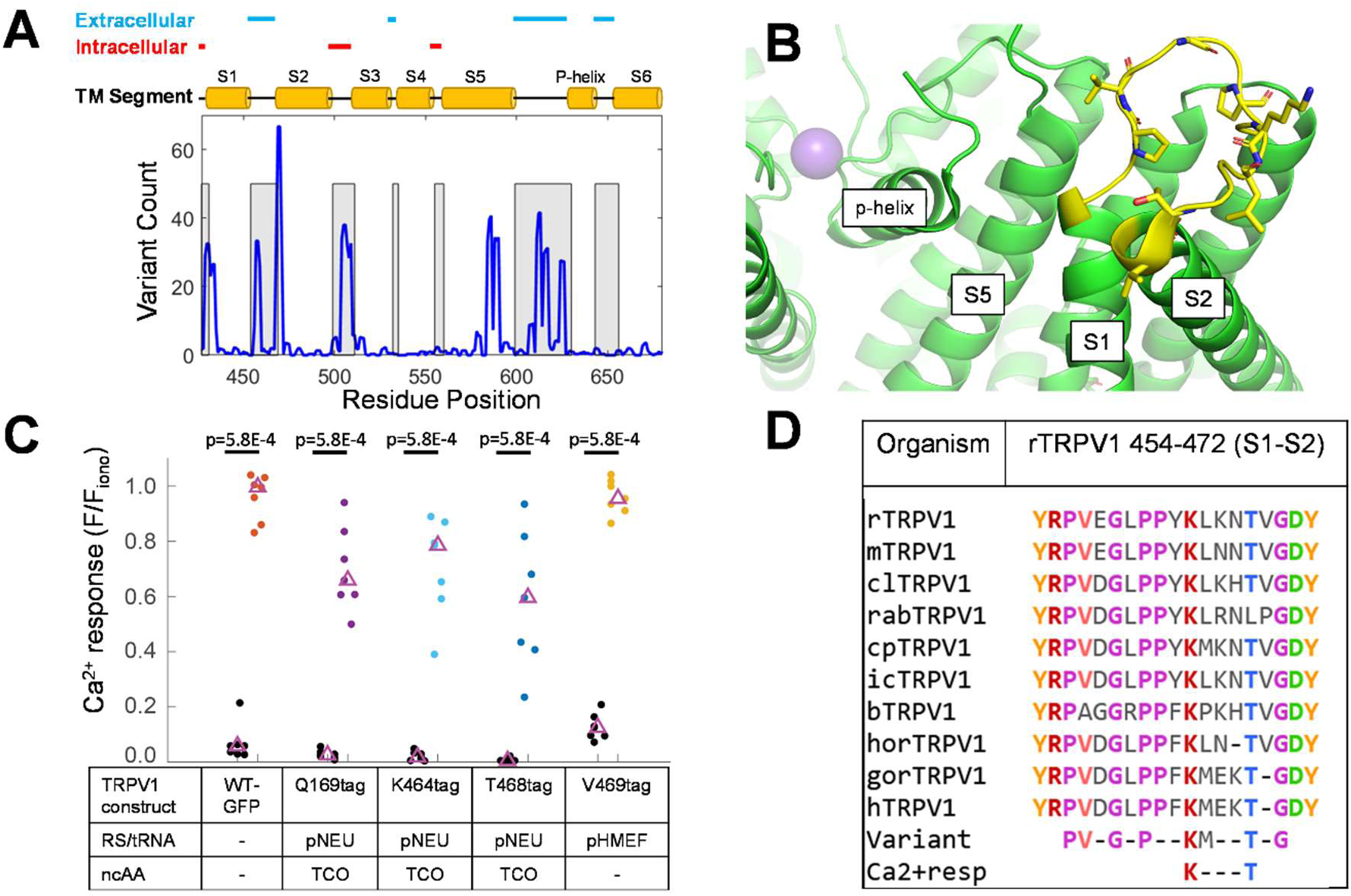
Insertion of ncAAs between transmembrane segments 1 and 2 of TRPV1. (A) Missense variants encoded in the TM domains of human TRPV1 (residues 427-680) as annotated in the gnomAD ver. 2.1.1. Blue line is the absolute variant count for the given residue filtered across 3 residues with a boxcar kernel. TM segments 1-6 are illustrated above the plot (yellow cylinders) and connecting loops are shown as grey regions within the plot. Extracullular (blue) and intracellular (red) loop regions above TM segments are potential sites for ncAA insertion. (B) Reconstructed structural model of one TRPV1 subunit from Cryo-EM data (PDB 7l2h green) reveals details of the extracellular loop between transmembrane segments 1 and 2 (yellow). (C) Ca^2+^ imaging experiments to determine successful expression of full-length TRPV1 constructs containing “tag” codons in the S1/S2 extracellular loop. RNA-synthetase/tRNA plasmid and type of ncAA used is given at the foot of the plot. Median value of set is indicated with purple triangle. Significant differences between Ca^2+^ imaging responses from non-responding cells (black) and cells with responses to 500nM capsaicin (colored) within the same imaging experiment were determined by Mann-Whitney U test. (D) Alignment of rat TRPV1 (rTRPV1) sequence (454-472; S1/S2 extracellular loop) against homologous sequences from several mammals (m: mouse,; cl: dog; rab: rabbit; cp: guinea pig; ic: thirteen lined ground squirrel; b: bovine; hor: equine; gor: gorilla; h: human). Variant positions identified in the gnomAD 2.1.1. database are listed below the conservation alignments with P456, V457, G459, P461, K464, T468, and G470 standing out as highly conserved residues (conservation >90%). Our first round of calcium imaging with P456tag, V457tag, K464tag, T468tag, and V469tag returned functional, ncAA-dependent responses with K464tag and T468tag.

### Electrophysiology

Currents were recorded at room temperature in symmetric divalent-free solutions (130 mM NaCl, 3 mM HEPES, 0.2 mM EDTA, pH 7.2) using fire polished borosilicate glass pipettes with filament (outer diameter 1.5 mm, inner diameter 0.86 mm; Sutter, MA). Pipettes were heat polished to a resistance of 3.5-7 MΩ using a micro forge. Cells were transfected with channel constructs and replated after 18-24 hours on 12 mm coverslips and placed in divalent-free solution in the chamber. 1 µM and 5 µM capsaicin solutions were prepared from a 5.3 mM capsaicin stock in ethanol. The same stock was diluted in the recording solution to 0.5 mM to make the 0.03 µM, 0.1 µM, and 0.3 µM capsaicin solutions. Excised inside-out patches were positioned at the opening of a tube in a “sewer pipe” configuration with 8 tubes connected to a rod controlled by an RSC-200 rapid solution changer (BioLogic, France). Capsaicin solutions (0.03, 0.1, 0.3, 1, and 5 µM) were perfused for 20-30s each as continuous 1s sweeps stepped the voltage from resting potential (10 ms) to -80 mV (100 ms), then up to +80 mV (300 ms) and back to resting potential (10ms). Currents were filtered at 5 kHz and recorded using an Axopatch 200A amplifier (Axon Instruments, Inc.) and PatchMaster software (HEKA). Data was analyzed in PatchMaster and IGOR (Wavemetrics, OR). Individual patch-clamp recordings with a complete set of stable current measurements in all capsaicin solutions were normalized to the maximal current with 5 µM capsaicin. Normalized data was fitted to a Hill equation to arrive at the K_1/2_ parameter: *I* = *I*_max_ ([ligand]*^n^*/(*K ^n^* + [ligand]*^n^*)), where n was permitted to float. Figure 4C reports the mean K_1/2_ for patch dose response measurements made with TRPV1 wild-type and TRPV1-exCellHalo. Statistical significance was determined by two-tailed Student’s t-test.

### Cell Culture

HEK293T/17 (ATCC: CRL-11268) cells were incubated in Dulbecco’s Modified Eagle Medium, supplemented with 10% fetal bovine serum, 50 μg <u>streptomycin</u>, and 50 units/mL penicillin, at 37°C and 5% CO2. Cells were passaged onto poly-L-lysine treated 25 mm coverslips. Cells were allowed at least 2h to settle onto the slips before being transfected using <u>Lipofectamine</u> 2000 (Life Technologies) as described in the manufacturer’s instructions. Cells transfected with TRPV1-tag mutants contained the target protein, TRPV1-tag-GFP in pcDNA3.1, an amber codon suppression plasmid, being either pNEU (ratio 1:1), pHMEF5-H1U6-RS (ratio 1:3), or pAcBac1-NES-R284 (ratio 2:1), and the media was supplemented with either 10uM LysZ (for pNEU), TCO (for pNEU or pHMEF5-H1U6-RS) or 30 μM Tetrazine3-Butyl (Tet3-Butyl for PAcBac1-NES-R284)(Kita et al., 2016; Serfling et al., 2019; Jang et al., 2020). Following transfection, Ca^2+^ imaging experiments were carried out after 24h, and western blot experiments were completed 48h after transfection.

### Molecular biology

The TRPV1-exCellHalo construct was built using the standard Gibson Assembly method of cloning (NEB, MA). Briefly, two primer sets were made for cpHaloTag in pCAG-HASAP (AddGene plasmid #138325) and rTRPV1: fragment set 5’-aggatgggaagaataactctgcctttgcccgcg-3’; 5’-ggtgtggactccataggcagcccggcaaattctggcc-3’ and vector set 5’-aatggccagaatttgccgggctgcctatggagtccacacca-3’; 5’-aaggtctcgcgggcaaaggcagagttattcttcccatcctcaatcagtg -3’. The TRPV1exCellHalo plasmid will be available through Addgene. All other TRPV1 constructs were made using either the Quickchange (Agilent, Santa Clara, CA) or in vitro assembly (IVA) methods of site-directed mutagenesis as previously described (Mott et al., 2023). For all mutant constructs, our wild-type TRPV1.pcDNA3 and TRPV1-cGFP.pcDNA3 vector was used as a template with either entirely (Quickchange) or partially (IVA) overlapping primers that contain the desired mutation. The full sequence of each completed construct was confirmed using Sanger sequencing.

### Ca^2+^ Imaging

Cells were grown on 25-mm coverslips and then incubated for 30 min at room temperature with Fluo-4 (AM; Thermofisher) at a concentration of 3 μM. The cells were then rinsed with Hepes buffered Ringer’s (HBR) solution (in mM, 140 NaCl, 4 KCl, 1.5 MgCl2, 5 D-glucose, 10 HEPES, and 1.8 CaCl2 and pH adjusted to 7.4 with NaOH) and allowed to rest for another 30 min in HBR at room temperature. The cells were imaged on a Nikon Eclipse Ti microscope using a 10× objective. For each slip, a brightfield image, a fluo-4 image, and a 3-min fluo-4 fluorescence movie with exposures of 100-ms and 0.5-s intervals were captured. During the time sequence, HBR is initially being perfused throughout the chamber. Perfusion is switched to 500 nM capsaicin in HBR at 30 s, and at the 2 min mark, 3 μM ionomycin is added to the chamber. The HBR and 500 nM capsaicin in HBR were perfused into the chamber using open, gravity-driven reservoirs, and 500 uL of 3 μM ionomycin was Pipetted into the chamber via micro-Pipette. The data obtained during these experiments were analyzed using Image J and Matlab (Mathworks, MA). For each time sequence, ROIs sized 10 × 10 pixels were placed over approximately seven responding cells where their level of fluorescence was tracked throughout the experiment. For each ROI, the proportion of the maximal fluorescence (F/F_iono_) that was reached with capsaicin (based on maximal response to ionomycin application) was calculated by removing a baseline offset. The Mann-Whitney U test was used to determine significance at and is given for each indicated comparison.

### DRG isolation and transfection with TRPV1exCellHalo

DRG isolation from wild type C57 mice was performed as described in Stein et al. (Stein et al., 2006). Additional supplementation of media with 100ng/ml nerve growth factor was done to maintain neuronal character as cells were kept for several days. Neuronal culture was transfected with TRPV1exCellHalo-pcDNA3.1 using JetPEI according to manufacturer’s instructions (Polyplus, FR). After 48 hours cells were incubated with 7μM Alexa660 HaloTag ligand in complete media for 15 minutes, which is twice the manufacturer’s recommended concentration (Promega, Madison WI). Sample was rinsed once with media and then incubated in complete media for an additional 20 minutes at 37 degrees C. Coverslips were imaged by epi-fluorescence with a 40x magnification objective in HBR.

### Live cell TRPV1 Imaging

Cells expressing either TRPV1-tag constructs or TRPV1exCellHalo were labeled with their respective click-chemistry fluorescent reagents in an imaging chamber during microscopy imaging experiments. TRPV1-tag pcDNA3.1 constructs were co-transfected into HEK293T/17 cells alongside pAcBac1-NES-R284 at a ratio of 1:2.in the presence of 30 μM Tet3-Butyl for 24-48 hours. Subsequent fluorescent labeling with 500 nM sTCO-Cy5 was done in the imaging chamber after exchanging cell culture media for HBR. Cells were incubated in HBR/sTCO-Cy5 labeling solution for 5 minutes followed by 3 minutes of continuous rinsing with HBR perfusion. Brightness correlation in Figure 3D and Figure S3-2 was done with 100X objective, GFP filter images captured via epi-fluorescence and near-TIRF field excitation of Cy5 dye with Cy5 filter set. Near TIRF means that the laser angle is set to illuminate the sample in an Epi/widefield configuration with reduced background signal because the light passes out to the side of the sample instead of transmitting directly out the top. The raw images shown in Figure S3-2 were assessed for mean GFP fluorescence in the indicated ROIs (1-5, mean value shown with each) and compared to maximal Cy5 fluorescence pixel in rectangular ROI for Cy5 channel (1-5, max pixel intensity reported with each). Rectangular ROIs were positioned such that smooth membrane outline fluorescence was captured and bright clusters were avoided. Sampling additional cells across different image fields was complicated by background offsets and strong dependence of intracellular GFP fluorescence signal on z-axis focal position, which would vary between image fields. Statistics are reported from a linear fit model (fitlm) function in Matlab. R-squared is the coefficient of determination and the p-value is returned from an F-test on the linear model with the reported data.

**Figure 3.**
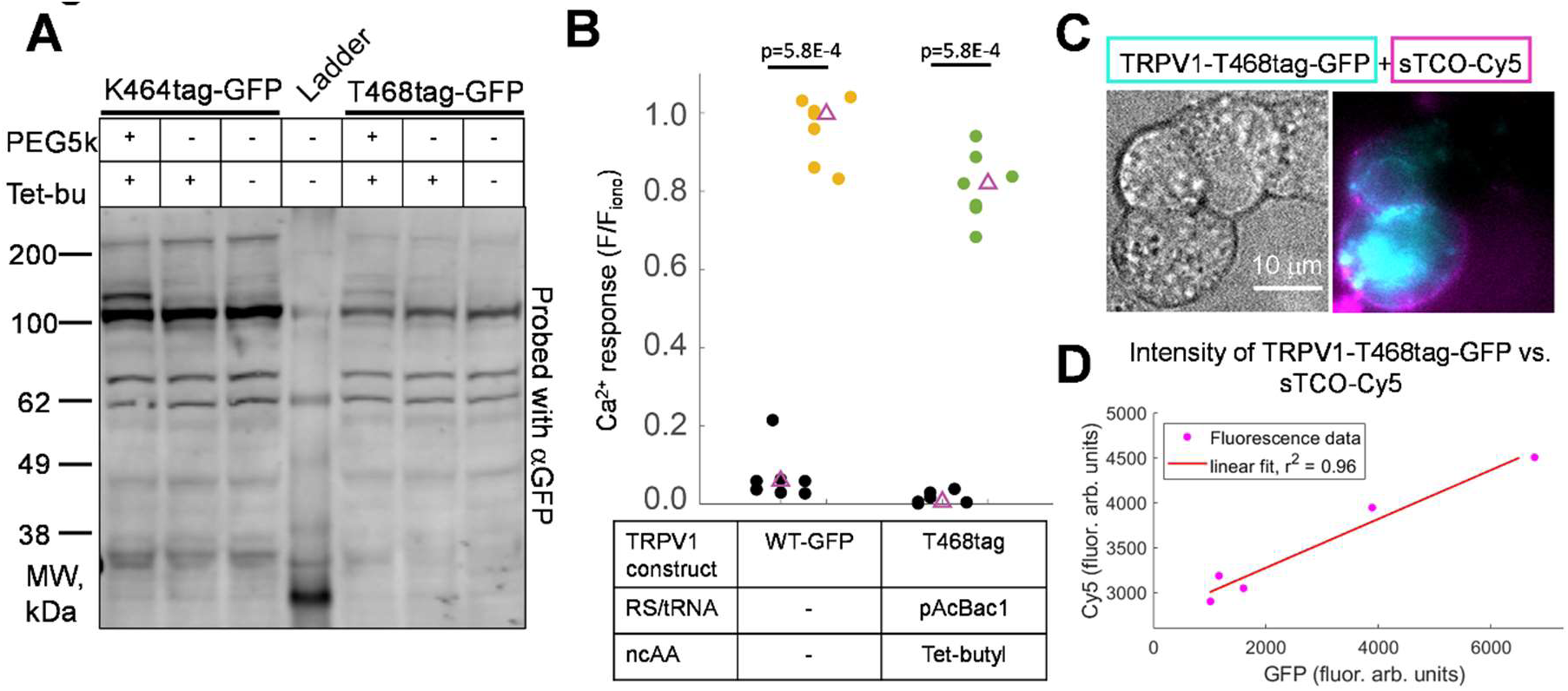
TRPV1-tag construct expression with GCE4All-Tetv3.0 system. (A) Western blot of TRPV1-K464tag-GFP and TRPV1-T468tag-GFP with pAcBac1 and Tet3-Butyl ncAA. Addition of sTCO-PEG5k (PEG5k) used to test access to tetrazine ncAA in TRPV1-tag target. Excess Tet3-Butyl (Tet-bu) is added to quench sTCO-PEG5k at the 10 minute point to prevent off target labeling with PEG5K. (B) Ca^2+^ imaging experiment to test functionality of TRPV1-T468tag-GFP expressed with GCE4All-Tetv3.0. Median value of set is indicated with purple triangle. Significant difference observed between Ca^2+^ response evoked by 500 nM capsaicin in TRPV1-T468tag-GFP cells (green) and non-responsive cells (black) in the same imaging experiment (Wilcoxon rank sum; p < 0.001). (C) HEK293T/17 cells expressing TRPV1-T468tag/Tet3-Bu-GFP ( cyan) are labeled with sTCO-Cy5 (magenta). (D) Correlation of GFP brightness in image of TRPV1-T468tag/Tet3-Bu-GFP expressing cells that are labeled with 500 nM sTCO-Cy5 for 5 minutes as described in methods. 5 cells expressing various levels of the TRPV1-T468tag/Tet3-Bu-GFP construct as determined by GFP intensity show correlated labeling with sTCO-Cy5 (linear fit, r^2^ = 0.96; p-value = 0.0034; see Figure S3-2 and Methods for description of statistics).

TRPV1exCellHalo was expressed from pcDNA3.1 in HEK293T/17 cells and cells were replated after 24 hours onto poly-D-lysine coated coverslips for imaging. After a minimum of 2 hours to allow for cell adhesion, labeling of TRPV1exCellHalo was completed in the imaging chamber by incubating cells with 3 μM Alexa660 Halo Ligand for 5 minutes (Promega, WI) in HBR followed by 3 minutes of continuous rinsing with HBR perfusion.

### Single molecule imaging of TRPV1exCellHalo

Expression of TRPV1exCellHalo in HEK293T/17 cells followed as previously described. After 24 hours, cells were transferred from 100mm dishes to coverslips treated with poly-D-lysine (p7886, Millipore-Sigma) to facilitate adhesion. 4-6 hours after replating the cells, we transferred the coverslips to our imaging chamber, labeled TRPV1exCellHalo, and imaged single channels with the following protocol:

1. Incubate cells in chamber with 2 μM Alexa488 HaloTag ligand in HBR for 5 minutes.
2. Rinse cells with continuous perfusion of HBR for 5 minutes.
3. With 100x objective and TIRF imaging locate cell with crowded, yet still resolvable single channels in cell footprint (smooth fluorescence in cell footprint indicates too high of expression)
4. Photobleach cell footprint with highest laser setting (our system is powered by a 50mW CW laser and we ordinarily maintain a neutral density filter in the laser path; this filter is removed and the laser operated at maximum power) for approximately 5 seconds. If the EM gain is lowered on the camera, the photobleaching process can be monitored to observe when adequate bleaching is reached.
5. Allow recovery of fluorescent TRPV1exCellHalo in the cell footprint for 5 minutes.
6. Acquire one 2.5 second movie (20Hz) with optimal gain (EM:300 for Andor iXon Life camera) for single channel imaging and laser intensity modulated to reduce bleaching (we place neutral density filter back into the path and power laser at 50%). This movie is to check if channels diffused sufficiently back into the footprint.
7. Begin acquisition of pre-capsaicin movie for 2.5 seconds (20Hz)
8. Rapidly deliver 5μM capsaicin solution in HBR into the chamber
9. Wait 15 seconds after the end of the pre-capsaicin movie to start recording the post-capsaicin movie with identical imaging parameters.

Mock treated samples follow an identical protocol with HBR added rapidly into the chamber instead of capsaicin solution. Because we wished to see what effect that plating conditions would have on the track displacements of TRPV1exCellHalo, we repeated the mock treatment protocol on cells that were plated on coverslips one day before the experiments. Displacement distributions from these mock treated cells generally exhibited higher mobility than their mock treated analogues plated on the same day as experiments (See Figure S5-1).

### Single molecule TRPV1 tracking

Data collected on moving TRPV1exCellHalo was processed using FIJI (https://imagej.net/software/fiji) by tracking the movement of ion channels within selected ROIs using TrackMate (v3.7.0)(Tinevez et al., 2017). Tracking analysis was performed on movies displaying channel activity just before the application of the capsaicin solution as well as movies displaying channel activity after the capsaicin solution was applied. To test the effects of the capsaicin, two movies were used on the same cell using a prior condition called pre-capsaicin and then a capsaicin solution added in the second.

1. Movies obtained from imaging were dragged and dropped into FIJI. Using the freehand selection tool, we selected an ROI over the first frame, taking care to avoid extremely bright clusters of spots in the movie while including as many individual spots as possible.
2. After applying the ROI to the movie, we used TrackMate, within FIJI, to track and analyze the movement of the channels throughout the movie. The TrackMate software was accessed from Tracking option within the drop down menu under the ’Plugins’ tab located on the horizontal taskbar across the top of the FIJI window.
3. The following settings and configurations were used when performing the tracking analysis using TrackMate:

1. Calibration settings were configured as follows: We set the pixel width and height to 0.160 um. The voxel depth and time interval was set to 1um and 0.052 seconds, respectively. Crop settings were set specific to the ROI of each movie by selecting the movie window and the clicking the ’Refresh source’ button on this page of TrackMate.
2. The LoG detector was selected to apply a Laplacian of Gaussian filter to the image.
3. The LoG detector settings were as follows: the estimated blob diameter to was set to 0.5 um and the threshold to 0.5. Additionally, sub-pixel localization was selected. *Clicking the Preview option on this page of TrackMate should display the selection of many fluorescent spots in the movie*.
4. This page described in detail the abilities and the statistics it will be able to perform.
5. The ’Initial Thresholding’ was set to restrict the sample of spots to be analyzed and tracked. We selected the ’Auto’ option on this page to consider the spots within the created Mask and chosen ROI.
6. The ’HyperStack Displayer’ View was selected.
7. No filters were applied to the spots. Additionally, ’Uniform color’ was selected as the color setting on this page.
8. The ’Simple LAP tracker’ was selected.
9. The Simple LAP Tracker was configured as follows: We specified the maximum linking and gap-closing distance to be 0.5 um. Additionally, the maximum gap-closing frame gap was set to 4 frames.
10. This page summarized the settings we selected to perform tracking.
11. No filters were applied to the tracks. The color of the tracks was set based on the ’Track displacement’.
12. The final page summarized the settings we chose for display options. Finally, we selected the ’Analysis’ button on this page to obtain the complete ’Tracking Statistics’ data set of the movie we examined.
4. All results from the ’Tracking Statistics’ sheet obtained from FIJI and TrackMate were selected and copied into an Excel spreadsheet for further analysis.

1. In Excel, the data was sorted in descending order based on column B, ’Number Spots’. We selected only the data with the value of ’Number Spots’ greater than 5 spots in order to only examine tracks of 5 spots or longer. The data of 5 number spots and higher was pasted in another sheet within the file.
5. We chose to analyze the specific tracking statistic, Track Displacement, using MATLAB (www.Mathworks.com, Natick, MA). Using MATLAB, we created figures representing trials with capsaicin or mock treatment in cells that were plated the same day or a day earlier.

1. For each experiment, we used MATLAB to create a figure displaying cumulative density histograms for the pre and post-capsaicin solution conditions. The Track Displacement data was imported into MATLAB and normalized from 0 to 1 in order to make the comparison between the different pre and post-capsaicin solution histograms possible. Both histograms were overlayed within one figure to visualize overall shifts in data.
2. We extracted the upper quartile values of both pre and post-capsaicin solution data sets. We used MATLAB to sort and index the data sets, before multiplying the data set by 0.75 and taking the floor value as the index number to obtain the upper quartile displacement value.
3. With two upper quartile values per experiment (before and after capsaicin). We plotted the pairwise comparisons in each experimental set.
4. Finally, in order to statistically analyze the potential effects of capsaicin activation, we performed a MATLAB two-sample Student’s t-test between the pre and post-capsaicin solution data sets for all experimental conditions.

### Western blotting

For western blotting, cells were incubated for 48 h at 37 C following transfection. The media was replaced after 24 h. Cells were collected from culture plates and washed with PBS before lysing with the following steps. The protein fraction was isolated by lysing the cells using Lysis buffer + HALT (25 mM Tris-HCL, 150 mM NaCl, 1 mM EDTA, 1% Triton, HALT [Thermofisher, MA]) and in accordance with the western blot protocol used by Zagotta and coworkers (Zagotta et al., 2016). The lysate was loaded into a PAGE SDS precast gel and was run at 200 V for 40 min. Using a BOLT wet transfer setup at 20 V for 60 min, the protein was transferred from the PAGE gel to a PVDF membrane (ThermoFisher, MA). The membrane was then pre-blocked in 5% skim milk in PBST for 1 h while shaking. The primary antibody (anti-GFP: TP401, Torrey pines Biolab, TX) was then added 1: 5000 directly to the blocking solution, and the membrane was covered and placed overnight on the shaker. The following morning, the membrane was rinsed with PBST and fresh 5% skim milk in PBST was added to the membrane. The secondary IR fluorescence labeled antibody was added 1:15000 to the blocking solution. Following a 45 min incubation in this solution while shaking, the membrane was imaged on a LI-COR imager with 700 nm and 800 nm wavelengths (LI-COR Biosciences, NE).

### PAGE gel-shift assay

Cell lysate from HEK293T/17 cells expressing the TRPV1-tag construct of interest with Tet3-Butyl as the ncAA was prepared as described in the previous section. Expression of sfGFP-N150tag with Tet3-Butyl was done by cotransfecting HEK293T/17 cells with pAcBac1-sfGFP-N150tag and pAcBac1-NES-R284 at an 8:1 ratio and supplementing the cell culture media with 30 μM Tet3-Butyl. Cell lysis buffer with 1% TritonX100 was used as described for Western blotting in Figure S3-1A. To each “+ sTCO-PEG5K” sample we added sTCO-PEG-5K for a final concentration of 100 μM. The reaction was quenched after 10 minutes or variable time by adding Tet3-Butyl to a final concentration of 3 mM. Treated lysate samples were loaded onto a PAGE precast gel and run at 200 V for 40 minutes. RIPA buffer was used to lyse cells for the gel shift assay in Figure S3-1B. To each “+ sTCO-PEG5K” sample we added sTCO-PEG-5K for a final concentration of 100 μM. The reaction was quenched after 10 minutes by adding Tet3-Butyl to a final concentration of 1.25 mM, for an additional 10 minutes. Laemmli buffer was added to each sample for a 1x concentration and then samples were loaded into a 12% SDS-PAGE gel and ran between 150 and 200 V, until the dye front ran off the gel. Western blotting to identify GFP bands was completed by first blocking with Licor blocking buffer. After this,

### Co-immunoprecipitation

Immunoprecipitation of TRPV1-cpHaloTag was done with Halo-Trap resin (catalog: ota) and as described in packaging instruction (Proteintech, IL) with the following changes: 1) For ∼6×10^6^ cells we used 500 μL lysis buffer of identical composition as used in the previously described western blot method. 2) Wash buffer contains in mM: 100 Tris pH 7.6, 150 NaCl, 0.1 EDTA, 1 mg/mL BSA, and 1x HALT. 3) Final wash step was done in wash buffer without BSA and HALT. 4) Elution for gel loading is in 40uL wash buffer and 1x SDS loading buffer (250 mM Tris pH 6.8, 8% SDS, 50% glycerol, 0.1% bromophenol blue), and 10% beta-mercaptoethanol. 5) Sample is not boiled prior to PAGE gel loading. Western blot analysis of TRPV1-cGFP pulldown fraction was done as previously described.

## Results

### Missense variants occur more frequently in intracellular and extracellular loops

We performed a bioinformatic analysis on human genomic data to replace an amino acid residue with an ncAA on the extracellular side of TRPV1, where it is unlikely to impact channel function. As the number of human genome sequences added to genomic variant databases increases, a growing body of research is applying this new-found sequence knowledge to protein structure and function (Plante et al., 2019; Bai et al., 2021). Previous studies of genomic missense variants occurring in the human population reveal a correlation pattern between buried side-chains in the TRPV1 N-terminus and a reduction in the number of variants for the corresponding residue (Mott et al., 2023). We have extended this pattern analysis into the transmembrane region, which contains conserved sequences assigned to the transmembrane segments, S1-S6, in TRPV1 and connecting segments of reduced conservation that correspond to internal and external loops (Figure 2A, *top*). We narrowed down potential sites for ncAA insertion in the external loops of TRPV1 by implementing an algorithm to count the absolute number of missense variants at each residue and plotted these data against the secondary structure of the transmembrane region (Figure 2A). The most salient feature of these plotted data is the increased occurrence of missense variants in regions connecting the transmembrane segments, precisely where we intend to replace an amino acid with an ncAA. However, the missense variant counts also show great variability within the extracellular and intracellular loops. Informed by the results of our bioinformatics study, we empirically determined the optimal placement of an ncAA, LysZ or TCO, at variant sites along the loop connecting S1 and S2, where the perturbation is expected to minimally affect the pore gating mechanism (Figure 2B, yellow). Success of the ncAA insertion in “tag” amber codon sites of TRPV1 was evaluated by Ca^2+^ imaging experiments in TRPV1-tag-GFP expressing HEK293T/17 cells. Previous work with TRPV1-Q169tag-YFP allowed us to evaluate TCO insertion with the pNEU amber codon suppression at novel sites in the S1-S2 loop (Zagotta et al., 2016). In comparison to background activity with cells that showed no appreciable change in fluorescence on exposure to 500 nM capsaicin, TRPV1-Q169tag expression in cells incubated with the TCO incurred a relatively significant response (Figure 2C, Q169tag black vs. purple markers). The difference in capsaicin response between transfected cells and background is shared by TRPV1 constructs with “tag” codons replacing the following amino acids in the S1-S2 loop: K464, T468, and V469 (Figure 2C). A notable result to our Ca^2+^ imaging experiments is the robust TRPV1-V469tag response obtained in the absence of an ncAA. Without co-expression of pNEU and supplementation with TCO there is no appreciable expression of full-length TRPV1-K464tag or TRPV1-T468tag, and this is in stark contrast to the expression product of TRPV1-V469tag visible in a western blot (Figure S2-1A). We investigated the properties of TRPV1-V469tag further by varying the circumstances under which the channel is expressed and determined that the TRPV1-V469tag-GFP construct requires neither co-expression of the pNEU plasmid or supplementation with an ncAA to achieve appreciable expression (Figures S2-1B and S2-2). Moreover, even in the presence of the suppression plasmid, pNEU, TRPV1-K464tag-GFP and TRPV1-T468tag still require the ncAA (TCO) to form functional channels in our Ca^2+^ imaging experiments (Figure S2-3). We summarize our results for scanning potential ncAA insertion sites in the S1-S2 loop in Figure 2D, which includes a sequence conservation alignment and illustrates the stepwise procedure we used from identifying missense variants in the human gnomAD sequence (“Variant") to experimentally assessing TRPV1-tag-ncAA-GFP functional channels.

### Cell surface fluorescence labeling of TRPV1 at single residues

The reduced expression level of previously characterized TRPV1-Q169tag-GFP construct with the pNEU support plasmid (Western Blot band in TRPV1-GFP lane compared to band in Q169tag-GFP of Figure S2-1) motivated us to test alternative amber codon suppression tools. We found that the tetrazine based ncAA insertion tools provided through the GCE4All Research Center was the optimal method for ncAA insertion and chemical modification of K464 and T468 (Jang et al., 2020; Ryan et al., 2022). In HEK293T/17 cells that express TRPV1 ‘tag’ mutations at K464 and T468, partial reactivity of the Tet3-Butyl ncAA with an sTCO-PEG5K substrate could be observed through an up-shift in the molecular weight bands of TRPV1-‘tag’-GFP (Figure 3A, lanes 1 and 5). Repeating the Ca^2+^ imaging study of TRPV1-T468tag-GFP with required GCE4All support plasmid (pAcBac1) and supplementation with ncAA Tet3-Butyl demonstrated a robust activation of channels in response to capsaicin (Figure 3B and Figure S3-1C). Animportant aspect of implementing amber codon suppression tools for surface labeling in pulse quench experiments is that the chemical reaction rate between the ncAA and chemical label must be rapid. We confirmed the rapid rate of labeling with the GCE4All system by expressing super-folder(sf) GFP as a ‘tag’ construct at position N150 in HEK293T/17 cells before lysing these and subjecting the free Tet3-sfGFP^150^ to pulsed labeling with sTCO-PEG5K for varying intervals of time. Our western blots of pulsed sTCO-PEG5K showed that maximal labeling of the Tet3-sfGFP^150^ is achieved in 3 minutes (Figure S3-1A, compare lane 2 (3 minutes) with lane 3 (10 minutes)). Our method for lysing HEK293T/17 cells and extracting TRPV1 proteins is, at least, partly the reason for why there is incomplete labeling of Tet3-sfGFP^150^ with sTCO-PEG5K. Under optimal sfGFP extraction conditions using RIPA buffer (see Methods) complete labeling of Tet3-sfGFP^150^ is possible with sTCO-PEG5K (Figure S3-1B). Ultimately, for the extracellular labeling of TRPV1-tag ncAA constructs to have any practical use reactivity with the sTCO-Cy5 dye needs to be sufficient for fluorescence imaging. To examine the threshold of reaction concentration and time needed for acceptable labeling of TRPV1-K464tag/T468tag-GFP channels, we expressed the construct in HEK293T/17 cells for 48 hours while in the presence of 30 μM Tet3-Butyl. After replating the cells on coverslips for imaging and mounting the coverslips in a chamber, we applied a labeling solution of 500 nM sTCO-Cy5 for 5 minutes before 3 minute of washing. Cells expressing TRPV1-T468tag-GFP as determined by GFP fluorescence were confirmed to have surface labeling by sTCO-Cy5 (Figure 3C). We were able to establish that the amount of sTCO-Cy5 label increased with expression of TRPV1-T468tag/Tet3-Bu-GFP by correlating the GFP fluorescence signal with Cy5 fluorescence on the cell surface (Figure 3D, Figure S3-2)TRPV1-K464tag/Tet3-Bu-GFP showed inconsistent sTCO-Cy5 labeling patterns (data not shown), and we elected to focus our continued efforts on the TRPV1-T468tag-GFP construct. Additional verification of the expression, function and labeling properties of TRPV1-T468tag/Tet3-Bu-GFP are reported in Koh and coworkers(Koh et al., 2024).

### Cell surface labeling of TRPV1 with circularly permutated HaloTag

Our amber codon suppression tools for labeling TRPV1 permit a small footprint labeling scheme with minimal perturbation to channel function. However, the amber codon suppression method for protein expression has much lower efficiency than a construct with the 20 naturally occurring amino acids (compare lane 1 and 2 of Figure S2-1A). To complement our single residue surface labeling approach with a high expression TRPV1 product, we engineered an insertion point for the circularly permutated (cp) version of HaloTag (Deo et al., 2021). Our TRPV1-exCellHalo construct contains a modified turret domain between transmembrane domain S5 and the pore helix with adequate space for the cpHaloTag to be accessible from the extracellular space in the fully assembled TRPV1 tetramer (Figure 4A). Ca^2+^ imaging experiments of TRPV1-exCellHalo showed robust responses to 500 nM capsaicin (Figure 4B). Evaluation of TRPV1-exCellHalo capsaicin sensitivity by dose response electrophysiology recordings revealed only a mild increased gain of function (Figure 4C). Apart from the non-significant increase in capsaicin sensitivity, TRPV1-exCellHalo appears to behave like the wild-type counterpart. To assess the labeling properties of the cpHaloTag domain on the extracellular side of TRPV1, we incubated HEK293T/17 cells that expressed the TRPV1-exCellHalo construct with 3.5 μM Alexa660-HaloTag for 5 minutes. Surface labeling with these conditions was sufficient for clear visualization of the plasma membrane on cells that also responded to capsaicin in a Ca^2+^ imaging experiment (Figure 4D). Since the TRPV1-exCellHalo construct was designed with the purpose of using it in cell culture and primary cell lines, we wished to test if the subunits containing cpHaloTag are able to co-assemble with intact TRPV1, containing a C-terminal GFP (TRPV1-cGFP). In an immunoprecipitation study of TRPV1-exCellHalo co-expressed with TRPV1-cGFP, we used a HaloTag antibody pulldown protocol and probed for TRPV1-cGFP with a GFP antibody. By pulling down the TRPV1-exCellHalo subunits of our co-expression lysate we obtained an enrichment of TRPV1-cGFP in our western blot, which was not the case for lysate derived from cells expressing only TRPV1-cGFP (Figures 4E and S4-1). Based on these results, we reason that exogenously expressed TRPV1-exCellHalo subunits may freely co-assemble with endogenously expressed TRPV1 in primary neurons.

**Figure 4.**
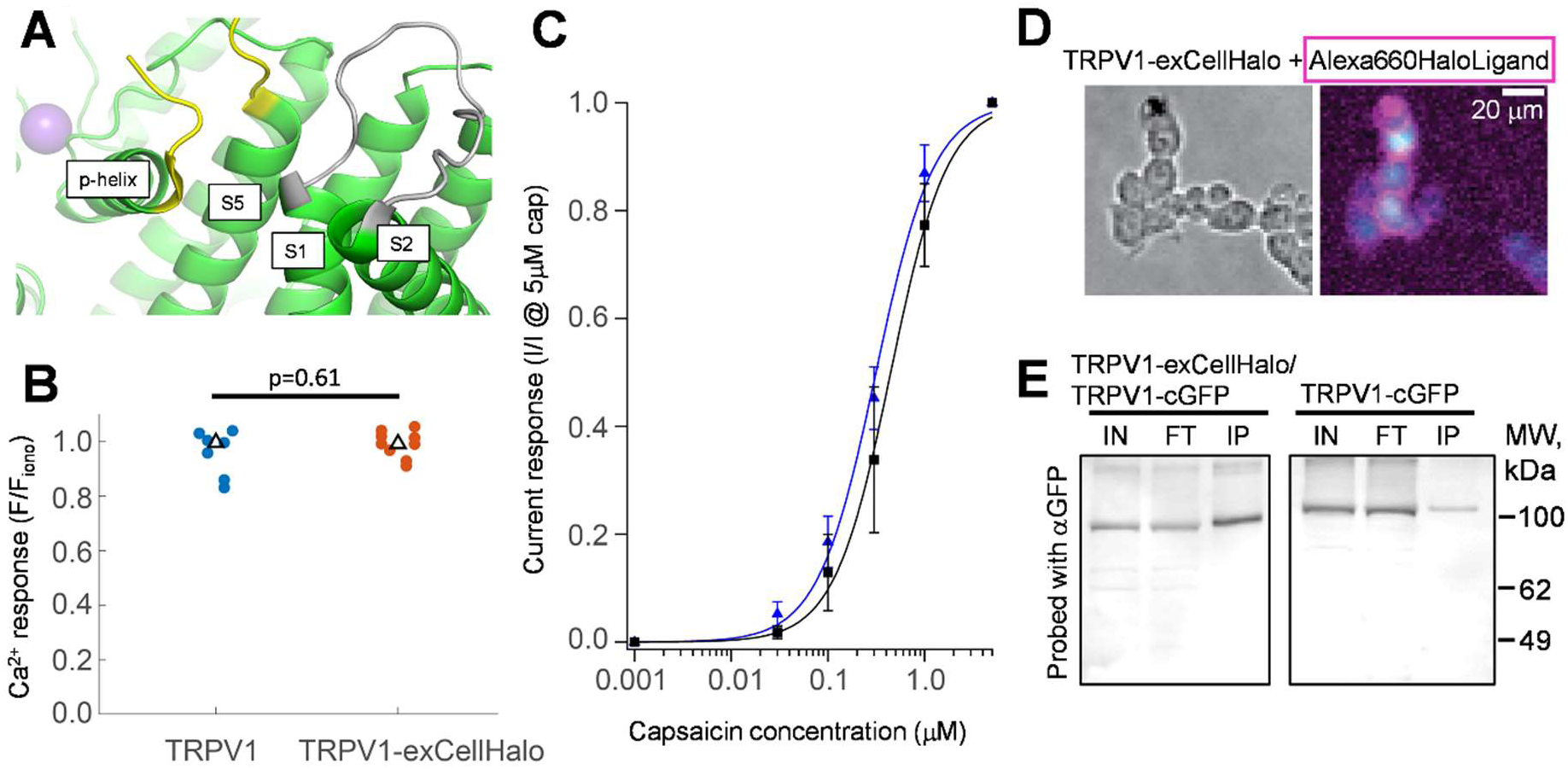
Expression and functional profile of TRPV1-exCellHalo. (A) Reconstructed model of extracellular region on a TRPV1 subunit (PDB 7l2h). Loop between TM S1 and S2 used for ncAA insertion is now shown in grey. Yellow termini demarcate the disordered region between TM segment 5 and the pore helix where circularly permutated HaloTag is inserted. (B) Ca^2+^ imaging experiment in HEK293T/17 cells comparing responses of wild-type TRPV1 against TRPV1-exCellHalo when activated by 500 nM capsaicin. Statistical comparison determined by Mann-Whitney U test. (C) Dose response experiment of wild-type TRPV1 (n = 4; K_1/2_ = 0.53 +/-0.28) and TRPV1-exCellHalo (n = 7; K_1/2_ = 0.34 +/-0.19). These K_1/2_ values are not significantly different as determined by Student’s two-tailed t test (p = 0.66). (D) DIC image (10x magnification; left) of HEK293T/17 cells transfected with TRPV1-exCellHalo. (Right) Fluorescence Ca^2+^-imaging experiment of TRPV1-exCellHalo expressing cells shown in left panel with Fluo4 following activation with 500 nM capsaicin (cyan). After Ca^2+^-imaging experiment, surface labeling is accomplished with incubation of HaloTag ligand Alexa660 (magenta) for 3 minutes. (E) Western blot of immune precipitation with anti-HaloTag antibody (Chemtek). Cell lysate is incubated with anti-HaloTag resin and blot is probed with anti-GFP antibody. Two conditions tested are co-expression of TRPV1-exCellHalo/TRPV1-cGFP or TRPV1-cGFP alone (IN: input; FT: flow through; IP: immunoprecipitated).

### Cell surface tracking of TRPV1exCellHalo

The prospect of using a rapid labeling, photostable, and exclusively surface labeled TRPV1 construct appealed to our group’s interest in tracking cell surface bound receptors. Our previous study on activity dependent mobility of cell surface TRPV1 relied on a genetically encoded GFP fused to an intracellular domain of the channel(Senning and Gordon, 2015). TRPV1exCellHalo expressed on the cell surface is exclusively labeled by cell impermeable dyes while the intracellular fraction is not modified, and the rapid labeling process ensures that labeled TRPV1exCellHalo remains on the surface during the experiment. We devised a similar experiment to our previous single molecule study of TRPV1-GFP described in Figure 6 of Senning et al. (2015), evaluating the changes to channel mobility after activation by capsaicin. Because our previous TIRF experiments with TRPV1-GFP could not guarantee a uniform channel population on the cell surface, we had to resort to an elaborate analysis of the channel dynamics. Here we are able to measure the displacement of individual TRPV1exCellHalo on the surface of HEK293T/17 cells before and after cells are treated with capsaicin and carry out a direct comparison between the distribution of displacements under the different conditions (Figure 5A, B). Total displacements of TRPV1exCellHalo tracks recorded in a 2.5 second movie prior to the rapid addition of 5μM capsaicin have a distinctly faster cumulative density histogram profile (blue) compared to the histogram of tracks after capsaicin treatment (red) and recorded 15 seconds later (Figure 5C). For each data set we compared the upper-quartile displacement value (0.75; dashed black line in Figure 5C) to demonstrate the changes in displacements occurring with capsaicin or mock treatments (Figure 5D and Figure S5-1). A notable aspect to the mock experiments done on the same day of plating was that pre- and post-treatment upper-quartile displacements were lower than in the capsaicin set of experiments, and we were able to see these values increase to those observed in capsaicin experiments by plating the cells on coverslips one day before the experiment. Regardless of what value ranges the displacements showed initially in our mock experiments, we did not observe the significant decrease in displacements observed in our capsaicin treated experiments. Our study of single molecule TRPV1exCellHalo mobility could, therefore, substantiate our earlier study that activation of TRPV1 channels by capsaicin decreases the lateral movement of the channels on the cell surface. To assess the surface expression properties of TRPV1exCellHalo in a more physiological preparation future studies will rely on expression of the channel in isolated DRG neurons, which we have shown is possible along with satisfactory labeling by HaloTag ligands (Figure 5E).

**Figure 5.**
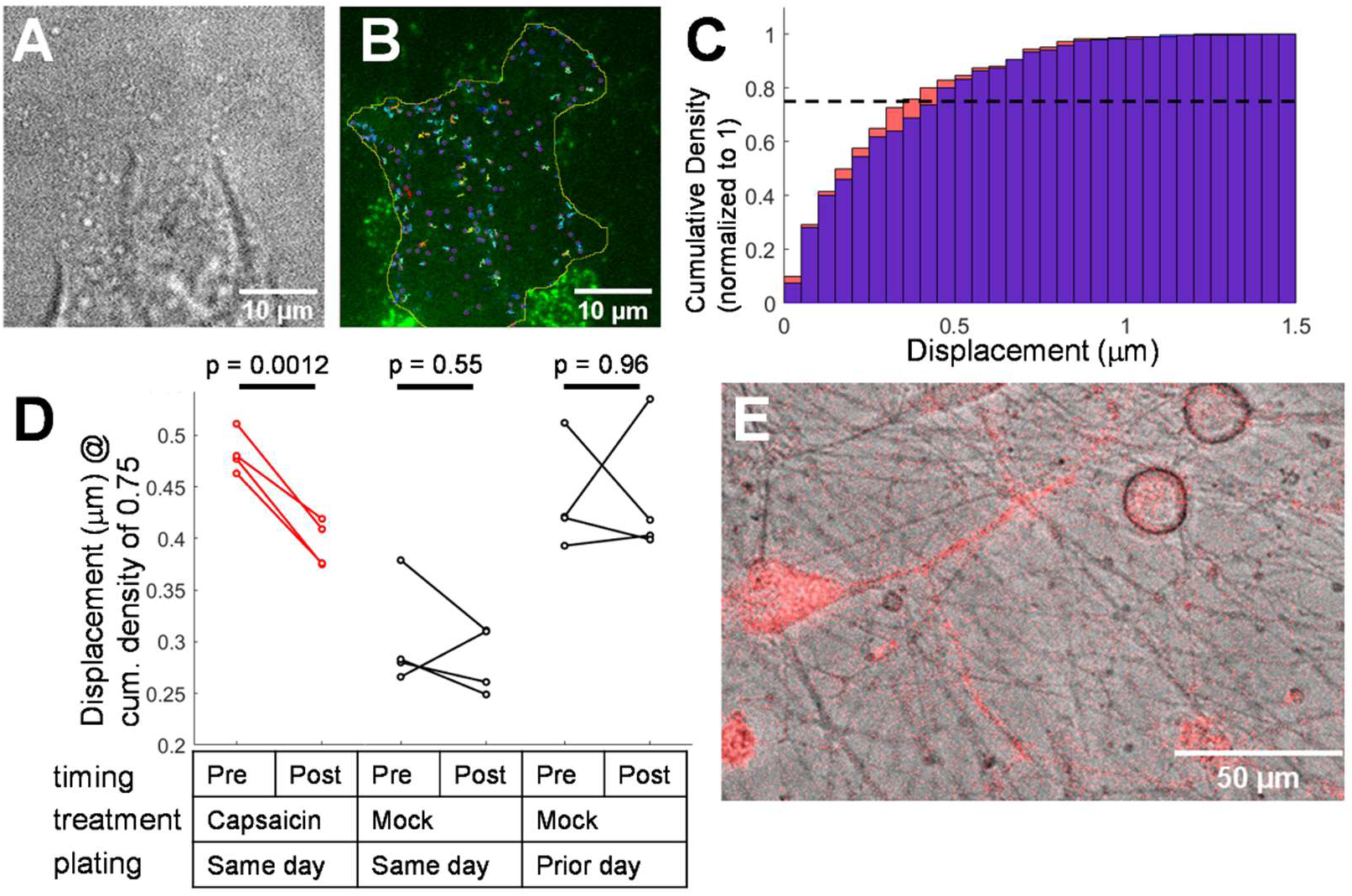
Reduced mobility of single TRPV1exCellHalo channels after capsaicin treatment. (A) DIC image of HEK293T/17 cells that transiently express TRPV1exCellHalo. (B) TIRF microscopy image of cell shown in panel A after labeling TRPV1exCellHalo with Alexa488 HaloTag ligand. The image shown is the first frame of a 2.5 second movie acquired pre-capsaicin treatment that reveals individual channels diffusing laterally across the cell surface (See Movie S1). Overlaid onto the image is the ROI (yellow) to define the area in which ion channels are tracked and individual tracks obtained during the movie. (C) Cumulative density histogram of individual channel track displacements. The total number of tracks is normalized to 1.0 in order to compare displacement distributions from tracks in a movie acquired pre-capsaicin (blue; tracks of panel B and Movie S1) to tracks acquired in a second (red) movie 15 seconds after rapid buffer exchange with a 5 μM capsaicin solution. Dashed, black line guides the eye to the upper-quartile (0.75) value in each distribution. (D) Quantitative comparison between track displacements from pre-capsaicin movies and post-capsaicin movies are made by evaluating the difference in the upper-quartile (0.75) displacement value (See full set of distributions in Figure S5-1). Mock treatment experiments with rapid buffer exchange that contains no capsaicin and day in which the cells are replated onto coverslips for experiment are noted in table. Statistical significance between pre- and post-capsaicin treatment is determined by paired Student’s t-test. (E) Transient expression of TRPV1exCellHalo in isolated mouse DRG culture followed by labeling with Alexa660 HaloTag Ligand.

## Discussion

There is a critical need to improve extracellular labeling methods for studies of channel trafficking and cell surface expression dynamics. Extensive use of labeling schemes ranging from immunofluorescent methods to FlaSH/ ReASH demonstrates the technological breadth of techniques used in these studies. However, experiments that examine protein dynamics are fundamentally constrained by the limitations imposed by the available labeling technology. For instance, in the case of immunofluorescent methods that target genetically encoded epitopes, concern arises due to the perturbation imposed by the large footprint of the antibody. On the other hand, the FlaSH and ReASH system, although sound in principle, is subject to off-target labeling, is sold as a cell permeant reagent, and has longer incubation times (∼30-60 minutes according to a manufacturer’s kit available through ThermoFisher) (Stroffekova et al., 2001; Specht et al., 2017). Many alternative strategies exist, and yet the benchmarks for technique optimization are unchanged: Use a minimally perturbing labeling scheme that is specific and rapid. We therefore sought to address these needs by setting two goals: 1) identify sites in extracellular regions of the ion channel TRPV1 that would tolerate perturbations from sequence substitutions and insertions without critically impacting function, 2) be able to implement pulse-chase time course experiments through a rapid labeling scheme. We also relied on a two-pronged strategy for labeling techniques to avert perceived or apparent weaknesses in one approach by exploring a complementary technique in parallel.

Our first approach implemented the amber codon suppression technique of replacing a single amino acid in the extracellular loop between S1 and S2, and we selected optimal ncAA insertion points by exploiting recently collected human genomic data available through gnomAD. We surveyed the extracellular loops, which connect transmembrane segments of the channel, and were able to refine our selection of ‘tag’ sites to experimentally test the feasibility of inserting the ncAA at a handful of positions between S1 and S2. Ca^2+^ imaging experiments used to test these sites revealed that K464, T468, and V469 could be substituted with the amber codon for protein termination, ‘tag’. However, results obtained under various expression conditions indicated that V469tag did not require the supplementary RNA synthetase and tRNA components for efficient translation of a full-length and functional TRPV1 channel. Although our experiments allowed us to eliminate potential ncAA insertion sites that were identified in our missense variant screen, we did not expect that V469tag would show such prominent readthrough, even when compared to supplementation with the pNEU plasmid and different ncAAs (Figure S2-1B). Translational readthrough at the ‘tag’ amber codon (+1, +2, and +3 positions) occurs as a sporadic event in eukaryotic cells and is partially determined by the flanking sequence nucleotides and interactions with the near-cognate tRNA or translational machinery (Dabrowski et al., 2015). Downstream nucleotide preference for a ‘c’ at the +4 position is a prominent marker for readthrough at stop codons ‘taa’ and ‘tga’, but for ‘tag’ a notable exception is made where multiple ‘g’s may appear at positions +4,+5, and +6 in gag-pol coding regions of mammalian type C retroviruses (Feng et al., 1992; Alam et al., 1999). In our TRPV1-V469tag sequence the nucelotides corresponding to the 2 codons after ‘tag’ are: ‘ggggac’. Perhaps more relevant to our task of inserting an ncAA, the Bultmann group devised an online tool (iPASS) to assess tolerance of flanking dna sequences to either side of a potential ‘tag’ site (Bartoschek et al., 2021). The iPASS tool returns a simple test score wherein a threshold over 1.0 indicates sufficiency for ncAA insertion. Both permissive sites we isolated, K464 and T468, returned scores of 1.16 and 0.87, respectively. However, the TRPV1-V469tag mutation and surrounding sequence gave an iPASS score of 0.05.

In a similar bid as implemented with ncAA incorporation, we attempted to insert the cpHaloTag domain at specific sites in the loop between S1 and S2 that coincided with increased missense variants. However, these constructs failed to express on the surface of HEK293T/17 cells. We could not readily discern the reason for this based on our experimental results, but a literature search for extracellular loop fusion proteins uncovered a recent study in which similar placement of genetically encoded fluorescent proteins in the ABCC6 integral membrane protein was also problematic, possibly due to short extracellular loops that would lead to folding or trafficking deficits if large polypeptide tracts were introduced (Szeri et al., 2021). Based on unsuccessful attempts at finding an insertion site between the S1 and S2 of TRPV1, we sought an alternate location and added the cpHaloTag in the “turret domain” of TRPV1 between S5 and the pore-helix (Cui et al., 2012). Concerns over whether a fusion protein above the pore domain would impact function were laid to rest when our Ca^2+^ imaging assays and e-phys dose-response experiments of the TRPV1exCellHalo construct nearly overlapped with wild-type function.

Once the ncAA insertion site was successfully determined by testing TRPV1 functionality, we addressed the second goal of our project to ask how we would improve the click-chemistry reaction rate between the ncAA and ligand reporter, such as a fluorescent dye molecule. Our experience with the pNEU vector using TCO as the ncAA was complicated by inconsistent yield during the labeling stage with the tetrazine-fluorophore. To address the erratic properties of the pNEU/TCO system we switched to the GCE4All amber codon suppression system that relies on a tetrazine based ncAA and sTCO ligand (Jang et al., 2020). Both our in vitro labeling reaction with an sTCO-PEG5K ligand and on cell labeling with sTCO-Cy5 could be performed within 5 minutes, which is well within the time needed for pulse chase experiments that would permit measurement of membrane trafficking events (Figure 3). Moreover, the study by Koh and co-workers made extensive use of the TRPV1-T468tag construct in a trafficking study and characterized its functional properties as well(Koh et al., 2024).

The rapid ∼5 minute reaction rate of HaloTag with its chloroalkane substrate is a valued property of this genetically coded click-chemistry product (Jonker et al., 2020). However, it was not certain whether our fusion of the circularly permuted HaloTag in the TRPV1exCellHalo construct would retain labeling with fluorescent ligands. Accessibility of the substrate binding site had been a concern given the permutated sequence of cpHaloTag and a position in our TRPV1 construct that imposed a crowded configuration above the pore-domain. In our hands, we could label surface TRPV1exCellHalo with Alexa660-HaloLigand in 5 minutes, which fulfilled our criteria for a rapid labeling step. Although our fluorescent HaloLigand assay was successful, we were not able to successfully use the HaloLink resin product (Promega, WI) as a pull-down matrix in a co-immunoprecipitation assay for heteromeric assembly of TRPV1exCellHalo and TRPV1-cGFP (data not shown). An alternative co-immunoprecipitation assay with IgG specific to HaloTag was still able to recognize the cpHaloTag and could confirm our hypothesis that TRPV1exCellHalo will heteromerically assemble with TRPV1-cGFP and presumably also wild-type TRPV1 subunits, and we speculate that the enzymatic site in TRPV1exCellHalo is occluded from matrix in the HaloLink product.

With the successful insertion of cpHaloTag into TRPV1 and intact capsaicin functionality, we moved forward with an experiment to test whether the mobility of TRPV1exCellHalo is changed by channel activation. Our experiment is a reprise of our study done with TRPV1-GFP in Senning et al. (2015). However, we can now track the subset of channels only expressed on the cell surface without signal contamination from intracellular pools. Our results with TRPVexCellHalo and Alexa488 HaloTag ligand reaffirmed the results from our earlier study: Capsaicin activation of TRPV1exCellHalo in Ca^2+^ containing buffer causes a reduction in mobility of the channel. An interesting observation in our current study was that the TRPV1exCellHalo channels exhibited different degrees of mobility based on how much time had passed since the cells were replated from 100mm dishes to poly-D-lysine coated coverslips. Evidently, the substrate under the HEK293T/17 cells would appear to influence the overall properties of channel mobility. Although there may be variability observed in channel displacement properties of cells that are replated on the same day (capsaicin and mock treated trials replated same day), the displacement shifts to indicate greater mobility when cells remain on the same substrate for longer times (mock trial on day after re-plating). That TRPV1 mobility could be sensitive to the underlying substrate of the cell is worth investigating further, especially considering that we do not know if this mechanism is mediated by direct contact with the extracellular matrix or adhesion factors within the cell.

Taken together, TRPV1-‘tag’/ncAA and TRPV1exCellHalo constructs form a comprehensive set of tools to investigate trafficking properties and cell surface expression of TRPV1. With only a single residue in TRPV1 converted to an ncAA in the amber codon suppression construct TRPV1-tag, we minimize the labeling perturbation to the site of an amino acid side-chain. The trade-off for a minimal label is the limited expression efficiency of the construct. Our second construct, TRPV1exCellHalo, takes advantage of genetically encoding the HaloTag domain in an extracellular loop at the expense of a larger molecular perturbation, but this construct comes with distinct advantages in expression efficiency while maintaining rapid labeling and near wild-type function. It is also noteworthy that the intracellular domains of TRPV1exCellHalo are unchanged and should be capable of sustaining all intracellular interactions. Moreover, the effect on cellular transport and expression by including the extracellular domain in TRPV1 can be studied and contrasted against the minimally perturbed TRPV1-‘tag’/ncAA channel.

## Acknowledgments

We thank Dr. Sharona Gordon for the construct TRPV1-Q169tag-YFP and advice on the study. We thank Dr. Nobumasa Hino for sharing pHMEF5-H1U6-RS, and we thank Dr. Marcel Goldschen-Ohm for sharing pNEU. This study was funded in part by start up funds from the University of Texas at Austin (to E.N.S.) and National Science Foundation grant no. 2129209 (to E.N.S.). This study was additionally supported by the GCE4All Biomedical Technology Development and Dissemination Center supported by the National Institute of General Medical Science grant RM1-GM144227.

## References

Alam, S.L., N.M. Wills, J.A. Ingram, J.F. Atkins, and R.F. Gesteland. 1999. Structural studies of the RNA pseudoknot required for readthrough of the gag-termination codon of murine leukemia virus. J Mol Biol. 288:837–852.

Bai, D., J. Wang, T. Li, R. Chan, M. Atalla, R.C. Chen, M.T. Khazaneh, R.J. An, and P.B. Stathopulos. 2021. Differential Domain Distribution of gnomAD- and Disease-Linked Connexin Missense Variants. Int J Mol Sci. 22.

Bartoschek, M.D., E. Ugur, T.A. Nguyen, G. Rodschinka, M. Wierer, K. Lang, and S. Bultmann. 2021. Identification of permissive amber suppression sites for efficient non-canonical amino acid incorporation in mammalian cells. Nucleic Acids Res. 49:e62.

Bessa-Neto, D., G. Beliu, A. Kuhlemann, V. Pecoraro, S. Doose, N. Retailleau, N. Chevrier, D. Perrais, M. Sauer, and D. Choquet. 2021. Bioorthogonal labeling of transmembrane proteins with non-canonical amino acids unveils masked epitopes in live neurons. Nat Commun. 12:6715.

Bevan, S., and J. Szolcsányi. 1990. Sensory neuron-specific actions of capsaicin: mechanisms and applications. Trends Pharmacol Sci. 11:330–333.

Blizzard, R.J., T.E. Gibson, and R.A. Mehl. 2018. Site-Specific Protein Labeling with Tetrazine Amino Acids. Methods Mol Biol. 1728:201–217.

Caterina, M.J., M.A. Schumacher, M. Tominaga, T.A. Rosen, J.D. Levine, and D. Julius. 1997. The capsaicin receptor: a heat-activated ion channel in the pain pathway. Nature. 389:816–824.

Chuang, H.H., and S. Lin. 2009. Oxidative challenges sensitize the capsaicin receptor by covalent cysteine modification. Proc Natl Acad Sci U S A. 106:20097–20102.

Cui, Y., F. Yang, X. Cao, V. Yarov-Yarovoy, K. Wang, and J. Zheng. 2012. Selective disruption of high sensitivity heat activation but not capsaicin activation of TRPV1 channels by pore turret mutations. J Gen Physiol. 139:273–283.

Dabrowski, M., Z. Bukowy-Bieryllo, and E. Zietkiewicz. 2015. Translational readthrough potential of natural termination codons in eucaryotes--The impact of RNA sequence. RNA Biol. 12:950–958.

Davis, J.B., J. Gray, M.J. Gunthorpe, J.P. Hatcher, P.T. Davey, P. Overend, M.H. Harries, J. Latcham, C. Clapham, K. Atkinson, S.A. Hughes, K. Rance, E. Grau, A.J. Harper, P.L. Pugh, D.C. Rogers, S. Bingham, A. Randall, and S.A. Sheardown. 2000. Vanilloid receptor-1 is essential for inflammatory thermal hyperalgesia. Nature. 405:183–187.

Deo, C., A.S. Abdelfattah, H.K. Bhargava, A.J. Berro, N. Falco, H. Farrants, B. Moeyaert, M. Chupanova, L.D. Lavis, and E.R. Schreiter. 2021. The HaloTag as a general scaffold for far-red tunable chemigenetic indicators. Nat Chem Biol. 17:718–723.

Feng, Y.X., H. Yuan, A. Rein, and J.G. Levin. 1992. Bipartite signal for read-through suppression in murine leukemia virus mRNA: an eight-nucleotide purine-rich sequence immediately downstream of the gag termination codon followed by an RNA pseudoknot. J Virol. 66:5127–5132.

Jang, H.S., S. Jana, R.J. Blizzard, J.C. Meeuwsen, and R.A. Mehl. 2020. Access to Faster Eukaryotic Cell Labeling with Encoded Tetrazine Amino Acids. J Am Chem Soc. 142:7245–7249.

Jonker, C.T.H., C. Deo, P.J. Zager, A.N. Tkachuk, A.M. Weinstein, E. Rodriguez-Boulan, L.D. Lavis, and R. Schreiner. 2020. Accurate measurement of fast endocytic recycling kinetics in real time. J Cell Sci. 133.

Karczewski, K.J., L.C. Francioli, G. Tiao, B.B. Cummings, J. Alföldi, Q. Wang, R.L. Collins, K.M. Laricchia, A. Ganna, D.P. Birnbaum, L.D. Gauthier, H. Brand, M. Solomonson, N.A. Watts, D. Rhodes, M. Singer-Berk, E.M. England, E.G. Seaby, J.A. Kosmicki, R.K. Walters, K. Tashman, Y. Farjoun, E. Banks, T. Poterba, A. Wang, C. Seed, N. Whiffin, J.X. Chong, K.E. Samocha, E. Pierce-Hoffman, Z. Zappala, A.H. O’Donnell-Luria, E.V. Minikel, B. Weisburd, M. Lek, J.S. Ware, C. Vittal, I.M. Armean, L. Bergelson, K. Cibulskis, K.M. Connolly, M. Covarrubias, S. Donnelly, S. Ferriera, S. Gabriel, J. Gentry, N. Gupta, T. Jeandet, D. Kaplan, C. Llanwarne, R. Munshi, S. Novod, N. Petrillo, D. Roazen, V. Ruano-Rubio, A. Saltzman, M. Schleicher, J. Soto, K. Tibbetts, C. Tolonen, G. Wade, M.E. Talkowski, B.M. Neale, M.J. Daly, D.G. MacArthur, and G.A.D. Consortium. 2020. The mutational constraint spectrum quantified from variation in 141,456 humans. Nature. 581:434–443.

Kita, A., N. Hino, S. Higashi, K. Hirota, R. Narumi, J. Adachi, K. Takafuji, K. Ishimoto, Y. Okada, K. Sakamoto, T. Tomonaga, S. Takashima, H. Mizuguchi, and T. Doi. 2016. Adenovirus vector-based incorporation of a photo-cross-linkable amino acid into proteins in human primary cells and cancerous cell lines. Sci Rep. 6:36946.

Koh, D.S., A. Stratiievska, S. Jana, S.C. Otto, T.M. Swanson, A. Nhim, S. Carlson, M. Raza, L.A. Naves, E.N. Senning, R. Mehl, and S.E. Gordon. 2024. Genetic code expansion, click chemistry, and light-activated PI3K reveal details of membrane protein trafficking downstream of receptor tyrosine kinases. bioRxiv.

Koplas, P.A., R.L. Rosenberg, and G.S. Oxford. 1997. The role of calcium in the desensitization of capsaicin responses in rat dorsal root ganglion neurons. J Neurosci. 17:3525–3537.

Leffler, A., B. Mönter, and M. Koltzenburg. 2006. The role of the capsaicin receptor TRPV1 and acid-sensing ion channels (ASICS) in proton sensitivity of subpopulations of primary nociceptive neurons in rats and mice. Neuroscience. 139:699–709.

Michaluk, P., and D.A. Rusakov. 2022. Monitoring cell membrane recycling dynamics of proteins using whole-cell fluorescence recovery after photobleaching of pH-sensitive genetic tags. Nat Protoc. 17:3056–3079.

Morenilla-Palao, C., R. Planells-Cases, N. García-Sanz, and A. Ferrer-Montiel. 2004. Regulated exocytosis contributes to protein kinase C potentiation of vanilloid receptor activity. The Journal of biological chemistry. 279:25665–25672.

Mott, T.M., J.S. Ibarra, N. Kandula, and E.N. Senning. 2023. Mutagenesis studies of TRPV1 subunit interfaces informed by genomic variant analysis. Biophys J. 122:322–332.

Nadezhdin, K.D., A. Neuberger, Y.A. Nikolaev, L.A. Murphy, E.O. Gracheva, S.N. Bagriantsev, and A.I. Sobolevsky. 2021. Extracellular cap domain is an essential component of the TRPV1 gating mechanism. Nat Commun. 12:2154.

Olah, Z., T. Szabo, L. Karai, C. Hough, R.D. Fields, R.M. Caudle, P.M. Blumberg, and M.J. Iadarola. 2001. Ligand-induced dynamic membrane changes and cell deletion conferred by vanilloid receptor 1. The Journal of biological chemistry. 276:11021–11030.

Plante, A.E., M.H. Lai, J. Lu, and A.L. Meredith. 2019. Effects of Single Nucleotide Polymorphisms in Human. Front Mol Neurosci. 12:285.

Ryan, A., O. Shade, A. Bardhan, A. Bartnik, and A. Deiters. 2022. Quantitative Analysis and Optimization of Site-Specific Protein Bioconjugation in Mammalian Cells. Bioconjug Chem. 33:2361–2369.

Senning, E.N., and S.E. Gordon. 2015. Activity and Ca²⁺ regulate the mobility of TRPV1 channels in the plasma membrane of sensory neurons. Elife. 4:e03819.

Serfling, R., L. Seidel, A. Bock, M.J. Lohse, P. Annibale, and I. Coin. 2019. Quantitative Single-Residue Bioorthogonal Labeling of G Protein-Coupled Receptors in Live Cells. ACS Chem Biol. 14:1141–1149.

Specht, E.A., E. Braselmann, and A.E. Palmer. 2017. A Critical and Comparative Review of Fluorescent Tools for Live-Cell Imaging. Annu Rev Physiol. 79:93–117.

Stein, A.T., C.a. Ufret-Vincenty, L. Hua, L.F. Santana, and S.E. Gordon. 2006. Phosphoinositide 3-kinase binds to TRPV1 and mediates NGF-stimulated TRPV1 trafficking to the plasma membrane. The Journal of general physiology. 128:509–522.

Stratiievska, A., S. Nelson, E.N. Senning, J.D. Lautz, S.E. Smith, and S.E. Gordon. 2018. Reciprocal regulation among TRPV1 channels and phosphoinositide 3-kinase in response to nerve growth factor. Elife. 7.

Stroffekova, K., C. Proenza, and K.G. Beam. 2001. The protein-labeling reagent FLASH-EDT2 binds not only to CCXXCC motifs but also non-specifically to endogenous cysteine-rich proteins. Pflugers Arch. 442:859–866.

Szeri, F., F. Niaziorimi, S. Donnelly, J. Orndorff, and K. van de Wetering. 2021. Generation of fully functional fluorescent fusion proteins to gain insights into ABCC6 biology. FEBS Lett. 595:799–810.

Szolcsányi, J., and Z. Sándor. 2012. Multisteric TRPV1 nocisensor: a target for analgesics. Trends Pharmacol Sci. 33:646–655.

Tinevez, J.Y., N. Perry, J. Schindelin, G.M. Hoopes, G.D. Reynolds, E. Laplantine, S.Y. Bednarek, S.L. Shorte, and K.W. Eliceiri. 2017. TrackMate: An open and extensible platform for single-particle tracking. Methods. 115:80–90.

Vetter, I., W. Cheng, M. Peiris, B.D. Wyse, S.J. Roberts-Thomson, J. Zheng, G.R. Monteith, and P.J. Cabot. 2008. Rapid, opioid-sensitive mechanisms involved in transient receptor potential vanilloid 1 sensitization. The Journal of biological chemistry. 283:19540–19550.

Wang, L., M.S. Frei, A. Salim, and K. Johnsson. 2019. Small-Molecule Fluorescent Probes for Live-Cell Super-Resolution Microscopy. J Am Chem Soc. 141:2770–2781.

Zagotta, W.N., M.T. Gordon, E.N. Senning, M.A. Munari, and S.E. Gordon. 2016. Measuring distances between TRPV1 and the plasma membrane using a noncanonical amino acid and transition metal ion FRET. J Gen Physiol. 147:201–216.

Zhang, X., J. Huang, and P.A. McNaughton. 2005. NGF rapidly increases membrane expression of TRPV1 heat-gated ion channels. EMBO J. 24:4211–4223.

