## Supplemental Information for "Fluorescence labeling strategies for the study of ion channel and receptor cell surface expression: A comprehensive toolkit for extracellular labeling of TRPV1"

Supplemental Figures

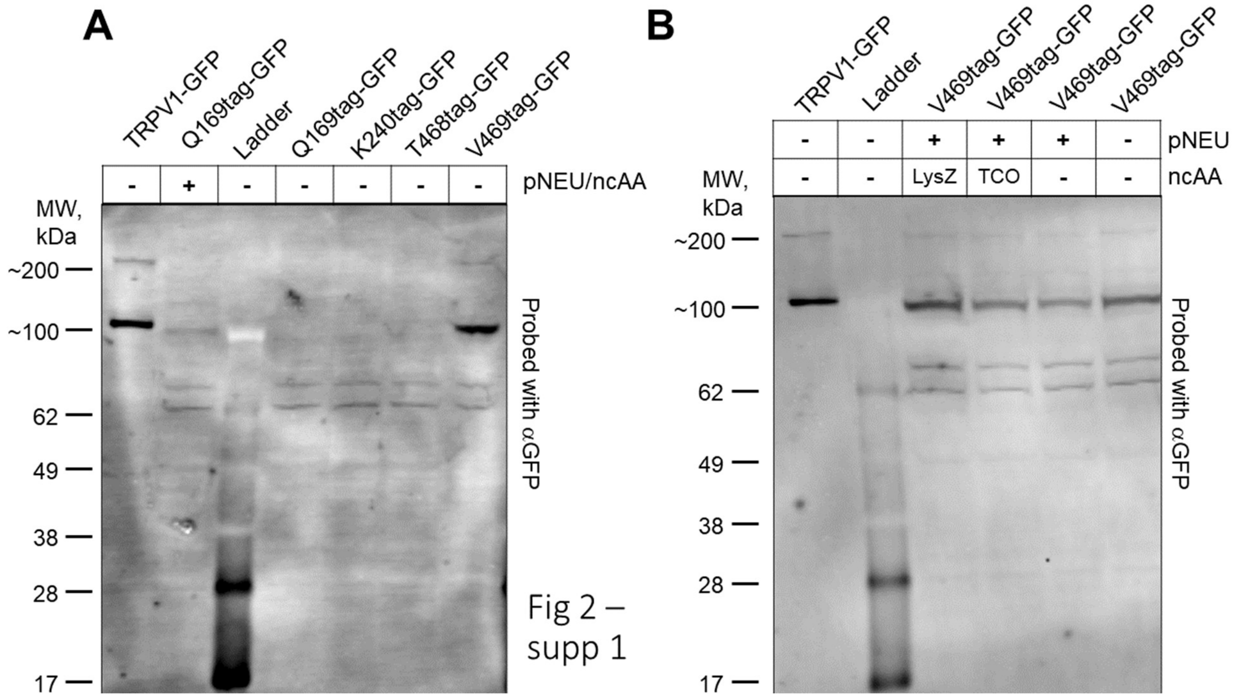


Figure S2-1. Protein expression of TRPV1-tag constructs. (A) Testing of construct TRPV1-Q169tag in the presence and absence of RNA synthetase/tRNA plasmid (pNEU) + ncAA (TCO). Lanes 5,6, and 7 contain expression products of TRPV1-K240tag, T468tag, and V469tag without pNEU. All lanes are probed with GFP antibody. (B) Testing of TRPV1-V469 in the presence and absence of pNEU and two different ncAAs (TCO or LysZ). Lanes are probed with GFP antibody.

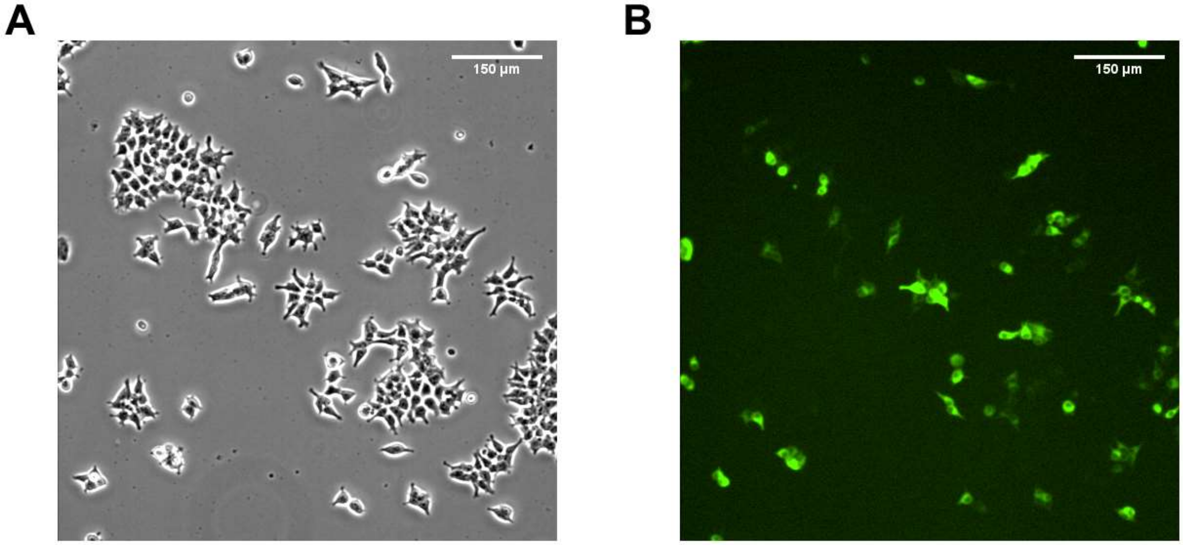


Fig 2 –
supp 2

Figure S2-2. Expression of TRPV1-V469tag-GFP in the absence of pyl-RNA synthetase/tRNA$^{pyl}$ (pNEU). (A) DIC image of HEK293T cells expressing TRPV1-V469tag-GFP. (B) Green fluorescence image of cells shown in panel A.

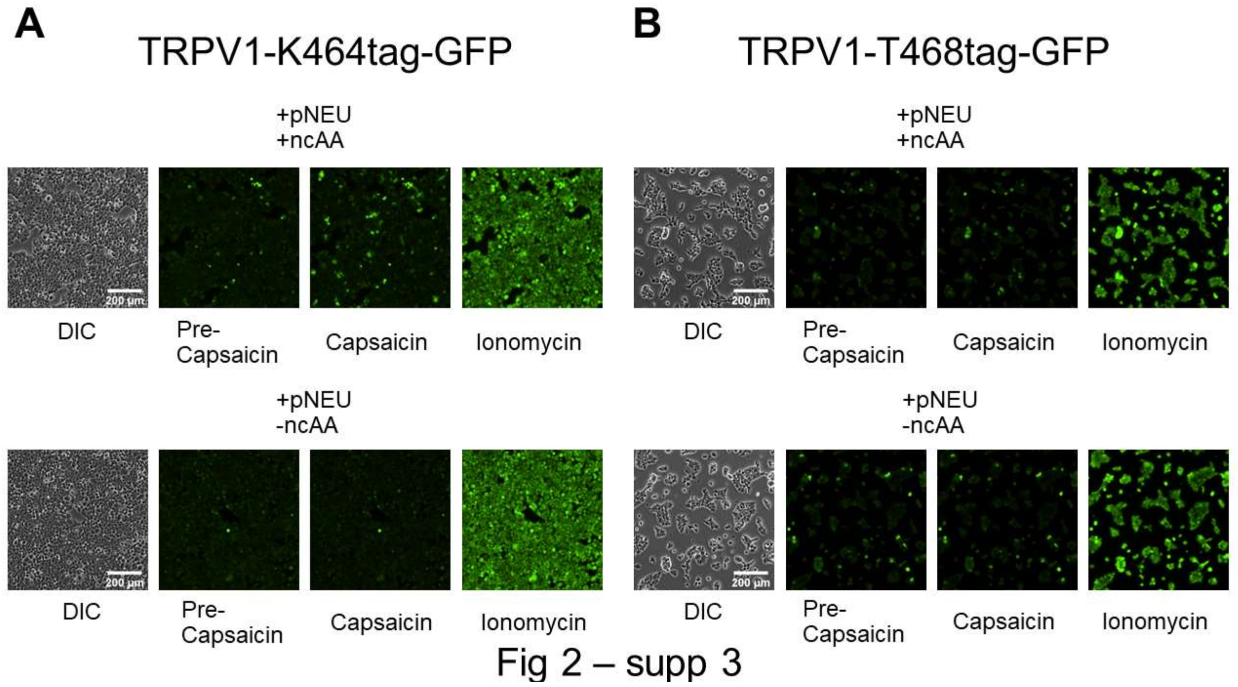


Figure S2-3. TRPV1-T468tag-GFP and TRPV1-K464tag-GFP negative control. Expression of functional TRPV1 channels is qualitatively absent when assessing Ca2+ imaging responses to 500nM Capsaicin without 30uM TCO (ncAA) in media. Neither TRPV1-T468tag-GFP (A) nor TRPV1-K464tag-GFP (B) generate Ca2+ responses in comparison to their transient transfection expression controls when supplemented with the pNEU plasmid but grown without TCO. All fluorescence experiment image sets are normalized to the brightness of their respective ionomycin image.

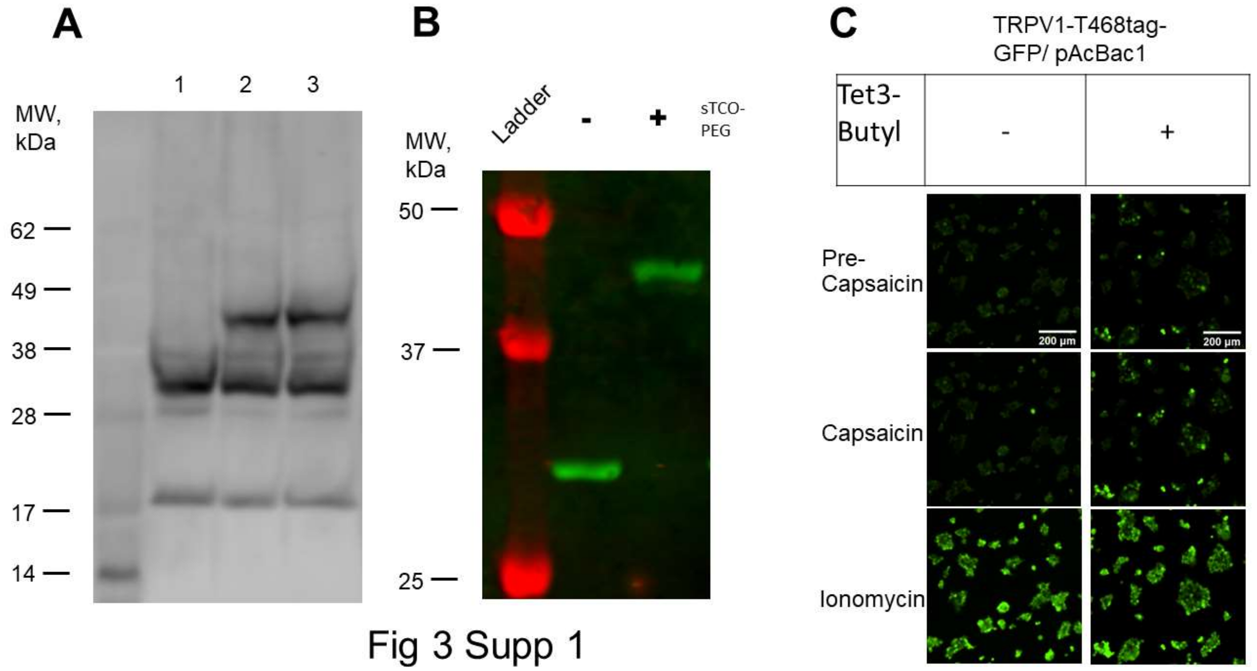


Fig 3 Supp 1

Figure S3-1. Western blots of gel-shift assays with sfGFP-N150tag/Tet3-Butyl and sTCO-PEG5K. (A) Extracts of sfGFP-N150tag/Tet3-Butyl expressed in HEK293T/17 pulse labeled with sTCO-PEG5K. Lane 1 represents sfGFP-150tag expressed in the presence of NES-R284 (pAcBac1) and Tet3-Butyl ncAA. Lane 2 demonstrates that addition of sTCO-PEG5k for 3 minutes induces a gel shift in the molecular weight band, and lane 3 shows the molecular weight shift of the sfGFP-N150tag after 10 minutes of incubation with sTCO-PEG5K. (B) Extraction of sfGFP-N150tag-FLAG/Tet3-Butyl with RIPA buffer and subsequent with (+) and without (-) sTCO-PEG5K labeling for 10 minutes before quenching with 1 mM Tet2 is shown after blotting for the C-terminal FLAG tag. (C) Control experiment of TRPV1-T468tag-GFP with pAcBac1 alone or supplemented with Tet3-Butyl in culture media. $Ca^{2+}$ imaging of TRPV1 activity done as described in methods using Fluo4-AM and 500nM capsaicin. All images are normalized to brightness of ionomycin image. Single responding cell in absence of Tet3-Butyl (-) was qualitatively deemed an outlier since cells otherwise showed no activity compared to Tet3-Butyl treated cells (+) which showed robust activity with capsaicin.

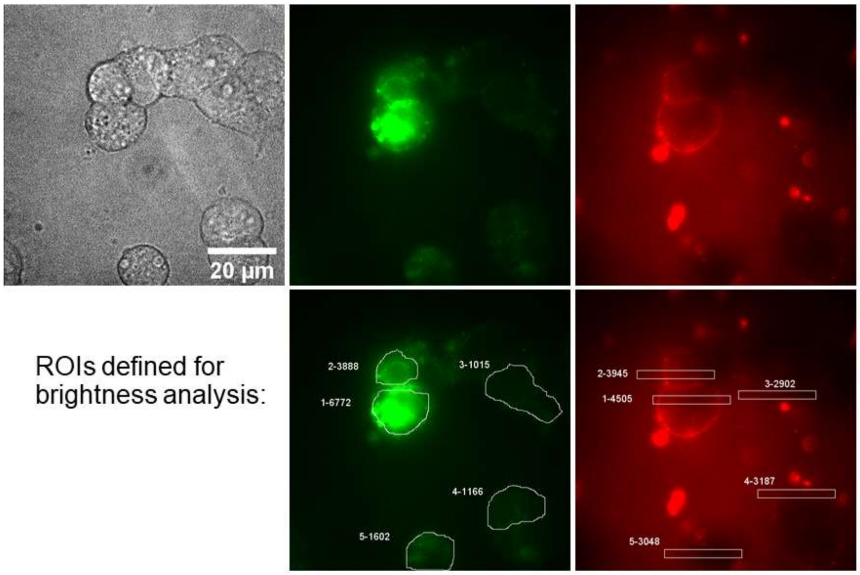


Fig 3
Supp 2

Figure S3-2. Brightness correlation analysis of TRPV1-T468tag/Tet3-Bu-GFP expressing cells and labeled with sTCO-Cy5. 100x objective image of HEK293T cells (DIC, top left) show varied degrees of GFP fluorescence from construct expression (GFP filter image, top middle) and correspondingly have different intensities of membrane labeling by sTCO-Cy5 (Cy5 filter image, top right). ROIs selected for mean GFP fluorescence in 5 cells (bottom middle) and Cy5 maximal pixel intensity (bottom right) are shown to illustrate our approach of capturing the full cell area to collect GFP fluorescence and selecting membrane regions of Cy5 that are representative of even, homogeneous dye distribution.

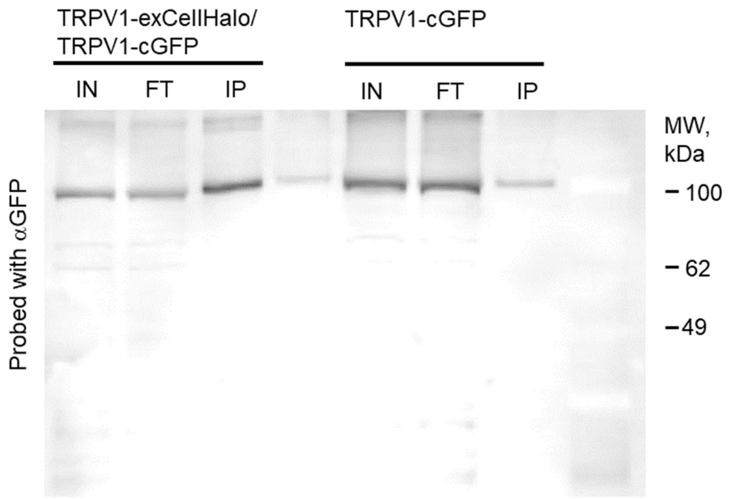


Fig 4
Supp 1

Figure S4-1. Western blot of anti-HaloTag immunoprecipitation. Blot is probed with anti-GFP. Lysate input (IN) and flow-thru (FT) from HEK293T/17 cells expressing either TRPV1-exCellHalo together with TRPV1-cGFP or TRPV1-cGFP run in lanes 1,2 and 5,6, respectively. Pull-down (IP) with anti-HaloTag is shown for different lysate samples in lanes 3 and 7. Ladder is in lane 8 but spectrally distinct from anti-GFP signal, which is why bands are faint.

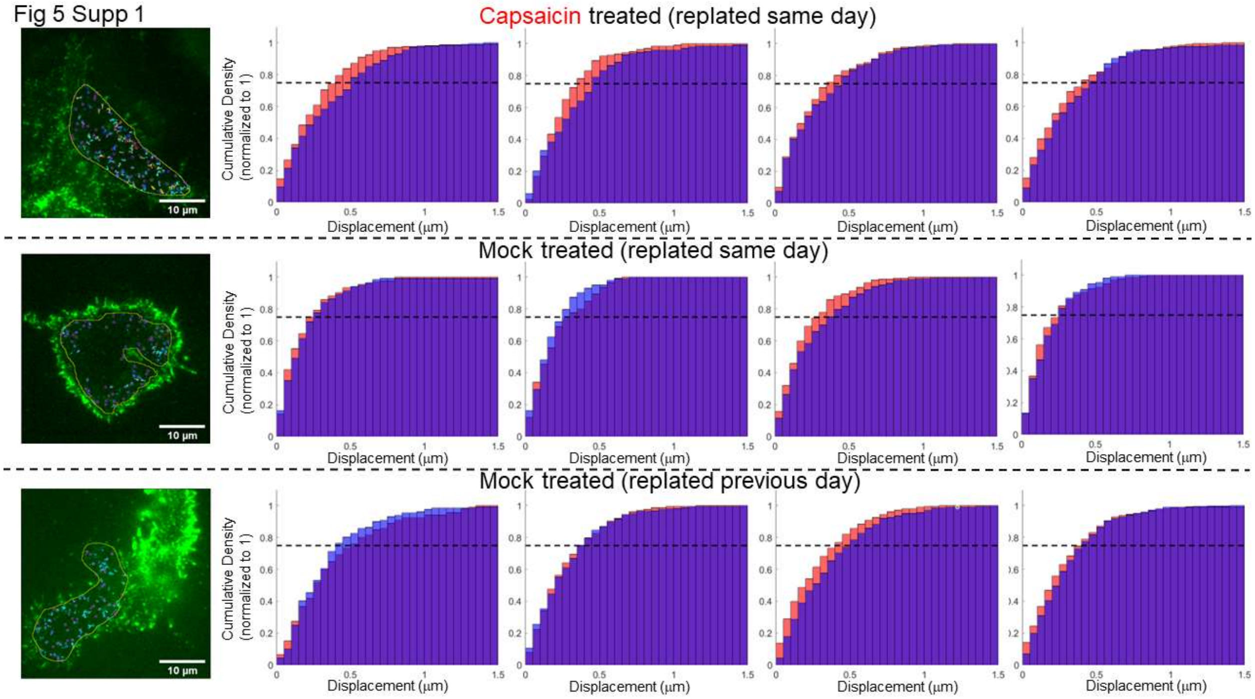


Figure S5-1. Cumulative density histograms of track displacements before and after capsaicin treatment. HEK293T cells expressing TRPV1exCellHalo are labeled with Alexa488 HaloTag ligand for single molecule tracking movies (See Methods). 2.5 second movies (50 frames) are acquired, 15 seconds apart, before and after 5μM capsaicin treatment to assess changes in TRPV1exCellHalo mobility. Cumulative density histograms (blue: pretreatment data; red: post-treatment data; purple: overlapping region in histograms) of track displacements are shown for three separate treatment conditions: (*top row*) Capsaicin treatment with TRPV1exCellHalo expressing cells that are replated the same day onto poly-D-lysine coated coverslips; (*middle row*) mock treatment of TRPV1exCellHalo expressing cells replated onto coverslips the same day; (*bottom row*) mock treatment of TRPV1exCellHalo expressing cells replated onto coverslips one day earlier. A guide-line at 0.75 is meant to help observe the upper-quartile values of the displacement distributions. Panels on the far left illustrate ROI bounded tracks from cells that are paired with data in the blue histogram in the adjacent panel. Summary plots for these data are shown in Figure 5D.

Supplemental Movies:

Movie S1. Image sequence capture under pre-capsaicin conditions in our capsaicin treatment sample. The first frame of the sequence is shown in Figure 5B and the analyzed track distances are displayed in the blue histogram of Figure 5C. Playback is at 0.5X speed.
